# Non-muscle myosin II acts non-cell autonomously to guide collective migration of the zebrafish polster cells

**DOI:** 10.64898/2026.06.15.727914

**Authors:** Amélie Elouin, Arthur Boutillon, Nicolas B. David

**Affiliations:** Laboratoire d’Optique et Biosciences, CNRS, Inserm, Ecole polytechnique, Institut Polytechnique de Paris, 91120 Palaiseau, France; Physics of Life, Technische Universität Dresden, Germany

**Keywords:** Non-muscle myosin II, collective cell migration, F-actin, protrusions, gastrulation, zebrafish

## Abstract

Collective cell migration is crucial in embryonic development and tumour dissemination. How cell-cell interactions guide collective cell migration is partially understood with a growing body of evidence pointing to a role for mechanical signals and intercellular tension. During zebrafish gastrulation, the anterior axial mesendoderm, also called polster, migrates collectively towards the animal pole. We here investigate the role of non-muscle myosin II (NMII), a well-known generator of intracellular tension, in this migration and uncover an unexpected non-cell-autonomous function. Inhibiting NMII activity, via genetic or pharmacological approaches, impairs both the speed and orientation of polster cells. Transplant experiments show that while NMII is required within the polster for proper cell orientation, it is not required in a cell-autonomous manner. Instead, NMII activity in neighbouring cells is essential to orient a given cell’s protrusions. Mechanistically, NMII localizes to the rear of cells and the base of actin-rich protrusions, where it promotes F-actin retrograde flow and generates mechanical tension. Using a conformation-specific α-catenin antibody, we show that NMII-dependent tension is required to open α-catenin in adjacent cells, providing a direct mechanotransductive link between neighbouring cells. These findings suggest a model in which protrusions not only serve as locomotory structures, but also as physical conduits for directional cues via transmission of a mechanical signal. Actomyosin-based contractility in one cell influences polarity in its neighbour, thereby ensuring directional coherence of the migrating group.

## Introduction

Cell migration is crucial both during development and pathological situations such as tumour dissemination and wound healing. Classic studies in cell culture show that cells move in a three-step manner; they extend their front through protrusions powered by Rac1 induced actin polymerization, attach to their substrate through adhesions, while RhoA activity in the body and rear resolves adhesion and contracts actomyosin to induce rear retraction (Yamada and Sixt, 2019). This model has been confirmed i*n vivo* for the migration of certain cell types, with a proposed grapple and pull model, in which cells extend a long protrusion, grip the substrate at the tip and pull on the protrusion to move forward (Fulga and Rørth, 2002).

*In vivo*, most cells move as part of larger cell groups, in so-called collective migrations, with the added complexity that, on top of propelling themselves forward, they need to coordinate their movements with neighbours (Friedl and Gilmour, 2009). One way to coordinate behaviours is through cell-cell interactions. One of the key findings over the past years was the identification of contact inhibition of locomotion (CIL) in which cells repolarize away from contact sites as a fundamental mechanism underlying collective motions in both mesenchymal and epithelial cells (Abercrombie and Heaysman, 1953; Carmona-Fontaine et al., 2008; Davis et al., 2015; Noordstra et al., 2023). Additional behaviour upon cell contacts, like contact following of locomotion or contact activation of locomotion, have been observed suggesting a wide range of responses to cell contacts and mechanisms to coordinate migration (Fujimori et al., 2019; Li and Wang, 2018). Another way to coordinate behaviours which emerged as a common theme in many collective systems is the creation of supracellular structures (Shellard and Mayor, 2019). In particular, the formation of actomyosin cables encircling the side and rear of the cell group have been observed in several systems, where they can both polarize the group, by restricting the formation of protrusions (Reffay et al., 2014; Wang et al., 2020) and induce coordinated cell movements through contraction (Pagès et al., 2022; Shellard et al., 2018).

We investigate how groups of cells coordinate their migration using the anterior axial mesendoderm of the zebrafish as a model. At the onset of gastrulation, the first cells to internalize on the dorsal side of the embryo are precursors of the polster (hereafter referred to as polster cells (Solnica-Krezel et al., 1995)). These cells form a group of approximately two hundred cells that migrate linearly from the embryonic organizer (the shield) towards the animal pole. Polster cells appear mesenchymal, with each cell in the group emitting actin-rich protrusions oriented in the direction of migration (Dumortier et al., 2012; Montero et al., 2003). Several pathways have been implicated in guiding polster migration, yet the nature of the instructive cues that direct this movement remains incompletely understood (Blanco et al., 2007; Čapek et al., 2019; Kai et al., 2008; Montero et al., 2005, 2003; Shimizu et al., 2005; Ulrich et al., 2005; Yamashita et al., 2004, 2002). We previously established that polster migration is a form of collective cell migration (Dumortier et al., 2012). E-cadherin–mediated cell–cell contacts are required for individual cells to orient their movement. More specifically, we recently showed that the mechanosensitive domain of α-catenin is required for cell-cell contacts to direct cell orientation (Boutillon et al., 2022). However, it is unclear what leads to the activation of this mechanosensitive domain. In epithelial cells, α-catenin acts as a mechanotransducer at adherens junctions, with its force-dependent unfolding leading to reinforcement of adhesion (Bazellières et al., 2015; Shewan et al., 2005). In such systems, non-muscle myosin II (NMII), acting on the junctional branched actin network, has been proposed as the force generator responsible for α-catenin opening (Heuzé et al., 2019; le Duc et al., 2010; Liu et al., 2010). We therefore asked whether NMII could similarly promote α-catenin opening and cell polarization in migrating polster cells, and set out to test the role of NMII in this process.

Using complementary approaches to inhibit NMII activity in polster cells during gastrulation, we found that NMII activity is required for effective polster cell migration, affecting both cell speed and orientation. Cell transplantation experiments revealed that NMII’s effect on migration speed is cell-autonomous, while its effect on orientation is surprisingly non-cell-autonomous. To investigate the basis of this non-cell-autonomous behaviour, we examined the subcellular distribution and function of NMII and found that it localizes to the base of protrusions, where it promotes F-actin retrograde flow and generates rearward tension. These findings led to a model in which NMII activity in one cell generates mechanical tension that opens α-catenin in the neighbouring cell, which we directly confirmed using a conformational α-catenin antibody. Together, these results uncover an unexpected non-cell autonomous role for NMII in generating intercellular forces that coordinate cell orientation. They also lead to revisit the function of protrusions, which not only serve as grapples to propel cells, but also function as mechanical transmitters of directional information that coordinate the cell group.

## Results

### NMII is required for polster migration towards the animal pole

To assess the potential role of NMII in polster cell migration, we targeted its upstream regulators, myosin light chain kinase (MLCK) and Rho-associated kinase (ROCK), which are primarily responsible for phosphorylating and activating the regulatory light chains of NMII (Lai et al., 2005; Totsukawa et al., 2000). Kinase dead versions of these enzymes have previously been shown to act as dominant-negative constructs and to effectively inhibit NMII activity in zebrafish (Amano et al., 2000, 1999; Blaser et al., 2006; Ishizaki et al., 1997; Marlow et al., 2002; Wadgaonkar et al., 2003). We injected mRNAs encoding dominant-negative MLCK (DN-MLCK) or ROCK (DN-ROCK) into one-cell stage *Tg(-1.8gsc:GFP)* embryos, allowing for clear identification of polster cells. *β-galactosidase* mRNAs were used as control, *H2B-mCherry* mRNAs were co-injected to label nuclei. At the doses used, embryos exhibited no overt morphological abnormalities during gastrulation (Fig. S1A). Nevertheless, immunostaining for phosphorylated myosin light chain (P-myosin) revealed a marked reduction in polster cells (Fig. S2), confirming successful inhibition of NMII activity. To quantify polster migration, we first measured the progression of the leading edge of the polster over a 40-minute period, starting at early gastrulation (60% epiboly). To ensure comparability across conditions, we verified that the leading edge was equidistant from the margin at the onset of measurement (Fig. S1B). Embryos expressing *DN-MLCK* or *DN-ROCK* exhibited a significant reduction in the progression of the leading edge of the polster (Figs. 1A, B and Movie S1). To determine whether this impaired progression towards the animal pole was due to reduced cell speed or impaired orientation, we tracked individual cells using labelled nuclei (Fig. 1C; Movie S1). Instantaneous speed and velocity in the direction of the animal pole (axial velocity) were measured (Figs. 1D, E). Instantaneous speed was moderately but significantly reduced in both *DN-MLCK* and *DN-ROCK* embryos (Fig. 1D; 21% and 14% reduction, respectively). Axial velocity was more strongly affected (Fig. 1E; 53% reduction in *DN-MLCK*, 37% in *DN-ROCK*). As reduction in instantaneous speed could not account for all the reduction in axial velocity, we looked at cell orientation, measuring the angle between each cell’s displacement vector and the direction of the animal pole (Fig. 1F). In control embryos, polster cell movements are strongly biased towards the animal pole, with 50% of displacement angles below 17°. In contrast, inhibition of NMII activity reduced cell orientation towards the animal pole: 50% of angles were below 35° and 32° in *DN-MLCK* and *DN-ROCK* embryos, respectively.

**Figure 1.**
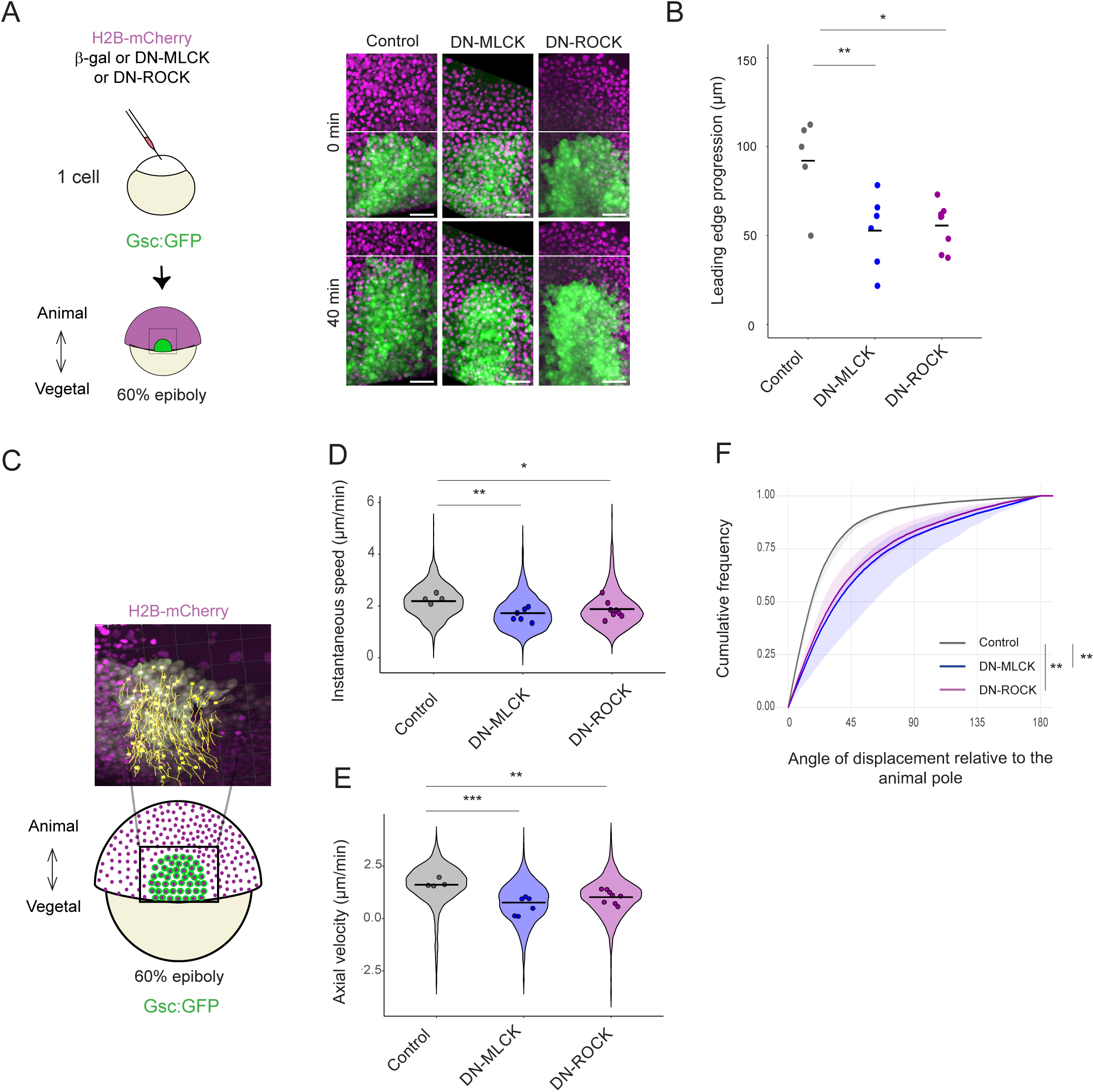
NMII is required for polster cell migration, contributing both to cell speed and orientation towar-ds the animal pole. (A) Representative dorsal views of *Tg(-1.8gsc:GFP)* embryos expressing *β-gal* (Control) or *DN-MLCK* or *DN-ROCK*, at early gastrulation (60% epiboly) and 40 minutes later. At t0, the white line marks the position of the leading edge of the polster in all three conditions; 40 minutes later, it marks its position in control embryos. Scale bar: 50 µm. (B) Leading-edge progression over a 40-minute period in control embryos (n = 5) and embryos ex-pressing *DN-MLCK* (n = 6) and *DN-ROCK* (n = 8). One-way ANOVA followed by Tukey’s HSD post-hoc tests. Adjusted p-values: Control vs *DN-MLCK*: 0.0097; Control vs *DN-ROCK*: 0.011. (C) Tracking of polster cells in *Tg(-1.8gsc:GFP)* embryos expressing *β-gal* (n = 1412 cells in 5 embryos) or *DN-MLCK* (n = 1891 cells in 6 em-bryos) or *DN-ROCK* (n = 1749 cells in 8 embryos). (D-F) Instantaneous speed, velocity towards the animal pole (axial velocity) and orientation of cell movement relative to the direction of the animal pole. (D, E) Circles on the violin plots indicate means per embryo. (F) The solid lines represent the overall Empirical Cumulative Distribution Function (ECDF) of angles for each condition, showing the proportion of data points with an angle less than or equal to a given value. The ribbons represent the interquartile range of the ECDFs calculated for each individual embryo within each condition. (D-F) Likelihood ratio test of a linear mixed effects model with treatment as a fixed effect and embryos as a random effect against a model without the fixed effect. Adjusted p-values: (D) Control vs *DN-MLCK*: 0.0028; Control vs *DN-ROCK*: 0.036. (E) Control vs *DN-MLCK*: 3.5E-4; Control vs *DN-ROCK*: 0.0013; (F) Control vs *DN-MLCK*: 0.0015; Control vs *DN-ROCK*: 0.0015.

NMII activity is therefore required in polster cell migration, contributing both to cell speed and orientation towards the animal pole. However, in these experiments, NMII was inhibited early in development, which could lead to other defects (Lai et al., 2005), secondarily responsible for the polster migration impairment.

### NMII is required during gastrulation for polster migration towards the animal pole

To temporally control NMII inhibition and assess whether its activity is specifically required during gastrulation for polster migration, we used pharmacological inhibitors: ML-7, which targets myosin light chain kinase (Lin et al., 2012; Odani et al., 2003), Y-27632, a Rho kinase inhibitor (Amano et al., 2000; Narumiya et al., 2000) and blebbistatin which slows down the release of phosphate after ATP hydrolysis, reducing the interaction of NMII with actin and thereby suppressing force generation (Kovács et al., 2004). Drugs were added slightly before the onset of gastrulation (germ ring stage). By mid-gastrulation, treated embryos showed no overt morphological abnormalities (Fig. S1D). Progression of the leading edge of the polster was measured over a 40-minute period, starting at early gastrulation (60% epiboly). As previously, we ensured that the leading edge was equidistant from the margin at the onset of measurement in all conditions (Fig. S1E). Embryos treated with ML-7, Y-27632 or blebbistatin all showed a reduction in leading edge progression (Figs. 2A, B and Movie S2). To investigate the cause of this reduced migration, individual polster cells were tracked in all conditions. Instantaneous speed and axial velocity (velocity in direction of the animal pole) were quantified as previously. Instantaneous speed was not significantly affected by ML-7, but was markedly reduced by blebbistatin (43%) and Y-27632 (24%) (Fig. 2C). Axial velocity was reduced in all three conditions (Fig. 2D; 37% reduction for ML-7, 49% for blebbistatin, 40% for Y-27632). Consistent with a more pronounced decrease in axial velocity compared to speed, cell orientation towards the animal pole was disrupted in all drug-treated embryos (Fig. 2E).

**Figure 2.**
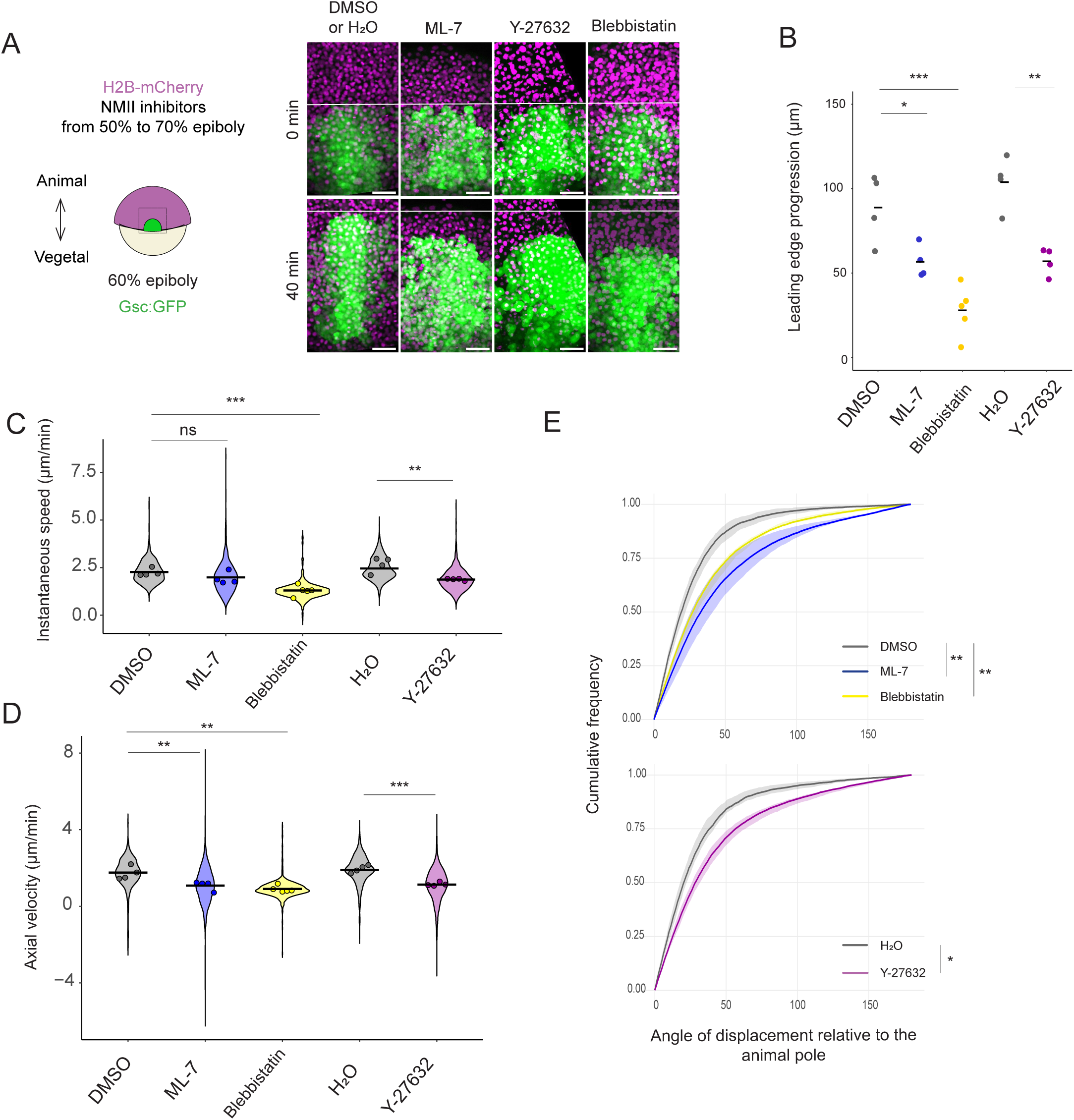
**NMII is required during gastrulation for polster cell migration.**(A) Representative dorsal views of *Tg(-1.8gsc:GFP)* embryos at early gastrulation (60% epiboly), treated with different NMII drug inhibitors, starting shortly before the onset of gastrulation (germ ring, 5 hpf). Embryos treated with DMSO or water only were used as control embryos. At t0, the white line marks the position of the leading edge of the polster in all conditions; 40 minutes later, it marks its position in control embryos. Scale bar: 50 µm.(B) Leading-edge progression over a 40-minute period in control embryos (n = 4, DMSO and H2O) and ML-7 (n = 4), Blebbistatin (n= 5) and Y-27632 (n = 4) treated embryos. One-way ANOVA followed by Tukey’s HSD post-hoc tests and t-test. Adjusted p-values: DMSO vs ML-7: 0.036; DMSO vs blebbistatin: 4.1E-4; H2O vs Y-27632: 4.3E- 3. (C-E) Instantatenous speed, velocity towards the animal pole (axial velocity) and orientation of cell movement relative to the direction of the animal pole. Circles on the violin plots indicate means per embryo. n = 517, 1360, 805, 630 and 1079 cells for DMSO, ML-7, Blebbistatin, H2O and Y-27632. Likelihood ratio test of a linear mixed effects model with treatment as a fixed effect and embryos as a random effect against a model without the fixed effect. Adjusted p-values: (C) DMSO vs ML-7: 0.089; DMSO vs blebbistatin: 1.4E-4; H2O vs Y-27632: 1.6E-3. (D) DMSO vs ML-7: 6.5E-3; DMSO vs blebbistatin: 6.1E-4; H2O vs Y-27632: 1.6E-5 (E) DMSO vs ML-7: 1.6E-3; DMSO vs blebbistatin: 1.6E-3; H2O vs Y-27632: 0.024.

These results demonstrate that NMII activity is required during gastrulation for effective polster cell migration toward the animal pole. However, gastrulation involves multiple tightly coordinated morphogenetic movements—including epiboly, internalization, convergence, and extension (Gong and Korzh, 2004) —which could also be affected by NMII inhibition. Previous studies have shown a key role for NMII in both epiboly and convergence/extension movements (Lai et al., 2005; Weiser et al., 2009). Thus, the impaired polster migration observed following drug treatments could arise from indirect effects on these other processes. Consistently, inhibition of NMII led to a broadening of the polster at the onset of measurement (Figs. S1C, F).

### NMII is required within the polster for cell orientation towards the animal pole

To assess whether NMII is required within the polster for its migration, we performed polster replacement experiments, in which the polster of wild-type embryos is replaced with polster cells expressing either *β-gal, DN-MLCK,* or *DN-ROCK* (Fig. 3A). This ensures that NMII inhibition is restricted to polster cells, while the rest of the embryo is wild-type. As in previous experiments, we ensured comparable initial conditions across embryos (Figs. S1G), and first monitored leading edge progression over a 40-minute period (Figs. 3A, B and Movie S3). Expression of either *DN-MLCK* or *DN-ROCK* in the grafted polster reduced its progression towards the animal pole. In contrast to whole-embryo NMII inhibition, tracking of individual polster cells revealed no significant change in instantaneous speed (Figs. 3C, D). Axial velocity, however, was reduced (Fig. 3E; 65% reduction for *DN-MLCK*, 37% for *DN-ROCK*). The uncoupling of speed and axial velocity pointed to a defect in orientation, which we confirmed by analysing the direction of cell displacements relative to the animal pole axis (Fig. 3F). These findings reveal that NMII activity is required within polster cells to ensure their proper orientation towards the animal pole.

**Figure 3.**
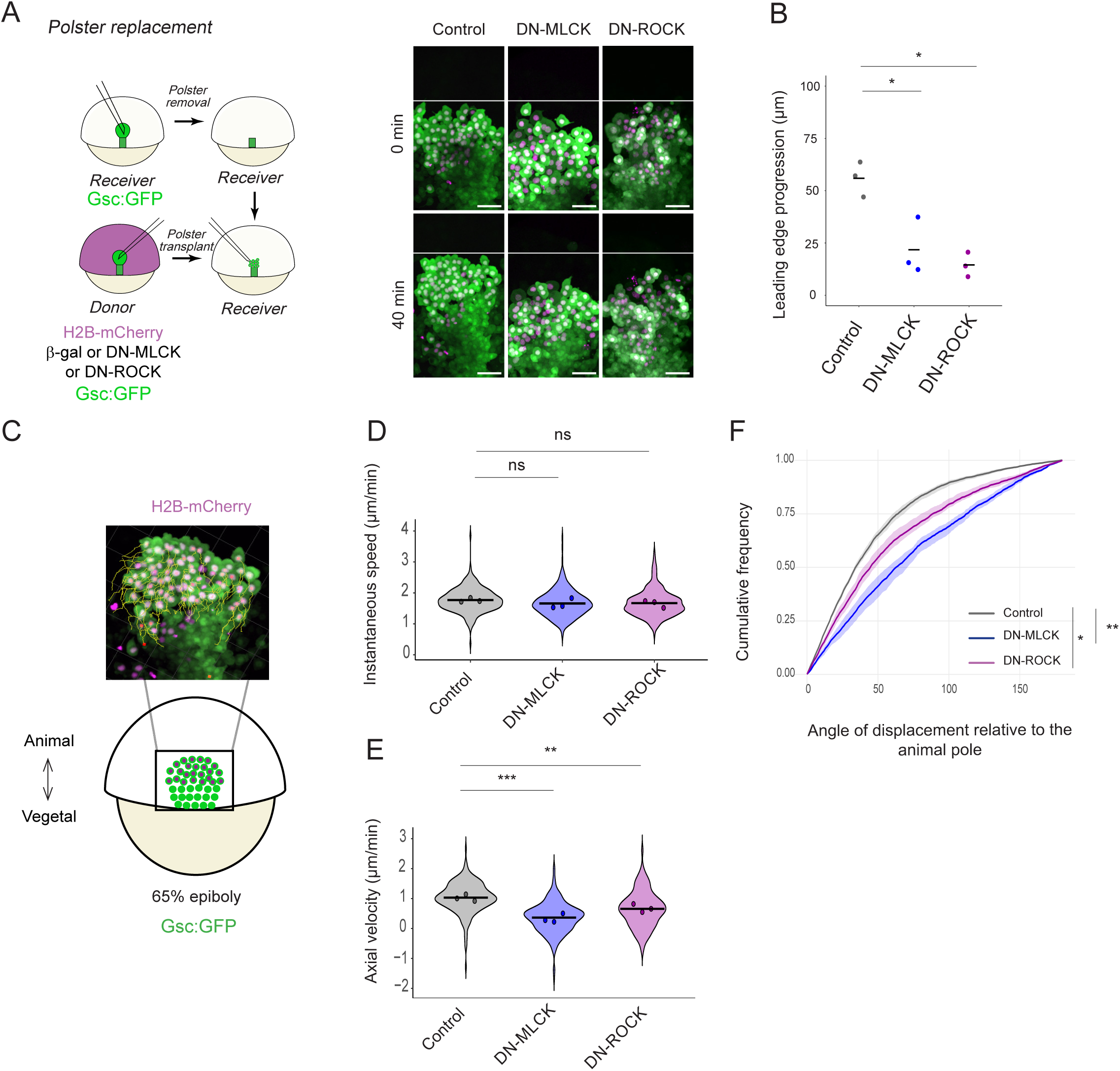
NMII is specifically required in the polster for its orientation towards the animal pole. (A) Representative dorsal views of *Tg(-1.8gsc:GFP)* embryos at early gastrulation (65% epiboly) in which the endogenous polster was replaced with polster cells expressing *β-gal* (Control) or *DN-MLCK* or *DN-ROCK*. At t0, the white line marks the position of the leading edge of the polster in all conditions; 40 minutes later, it marks its position in control embryos. Scale bar: 50 µm. (B) Leading-edge progression over a 40-minute period in embryos with polster cells expressing *β-gal* (control), *DN-MLCK* or *DN-ROCK*. n = 3 embryos for each condition. One-way ANOVA followed by Tukey’s HSD post-hoc tests, adjusted p-values: Control vs *DN-MLCK*: 0.013; Control vs *DN-ROCK*: 5.1E-3. (C) Tracking of polster cells expressing *β-gal* (n = 224 cells in 3 embryos) or *DN-MLCK* (n = 269 cells in 3 embryos) or *DN-ROCK* (n = 239 cells in 3 embryos). (D-F) Instantaneous speed, velocity towards the animal pole (axial velocity) and orientation of cell movement relative to the direction of the animal pole. Circles on the violin plots indicate means per embryo. Likelihood ratio test of a linear mixed effects model with treatment as a fixed effect and embryos as a random effect against a model without the fixed effect. Adjusted p-values: (D) Control vs *DN-MLCK*: 0.22; Control vs *DN-ROCK*: 0.22. (E) Control vs *DN-MLCK*: 3.0E-4; Control vs *DN-ROCK*: 4.7E-3; (F) Control vs *DN-MLCK*: 1.4E-3; Control vs *DN-ROCK*: 0.011.

### NMII is required non-cell autonomously in cell orientation towards the animal pole

To further investigate how NMII activity regulates polster cell orientation, we conducted cell transplant experiments to inhibit NMII in a cell-autonomous manner (Fig. 4A). We previously reported that individual poorly oriented—or even non-motile—cells, when transplanted into a wild-type polster, are passively carried along by neighbouring cells (Boutillon et al., 2022; Dumortier et al., 2012). As a result, their trajectories do not reliably reflect their intrinsic behaviour or orientation. However, their orientation can be assessed by examining the direction of their actin-rich protrusions. Wild-type polster cells frequently extend such protrusions, which are predominantly directed toward the animal pole (Dumortier et al., 2012; Figs. 4A, B, D). We therefore asked whether NMII activity regulates the dynamics and orientation of these protrusions. To address this, we transplanted single or very few actin labelled polster cells into the polster of wild-type host embryos (Fig. 4A). Inhibition of NMII activity via expression of *DN-MLCK* or *DN-ROCK*, reduced the frequency and increased the maximal length of actin- rich protrusions, revealing a role of NMII in controlling cell protrusiveness (Figs. 4B, C). Surprisingly, however, NMII inhibition had no effect on protrusion orientation, indicating that NMII is not cell-autonomously required for polster cell orientation (Fig. 4D). This absence of a cell-autonomous effect on cell orientation, while NMII is required within the polster for cell orientation, pointed to a potential non-cell-autonomous effect.

**Figure 4.**
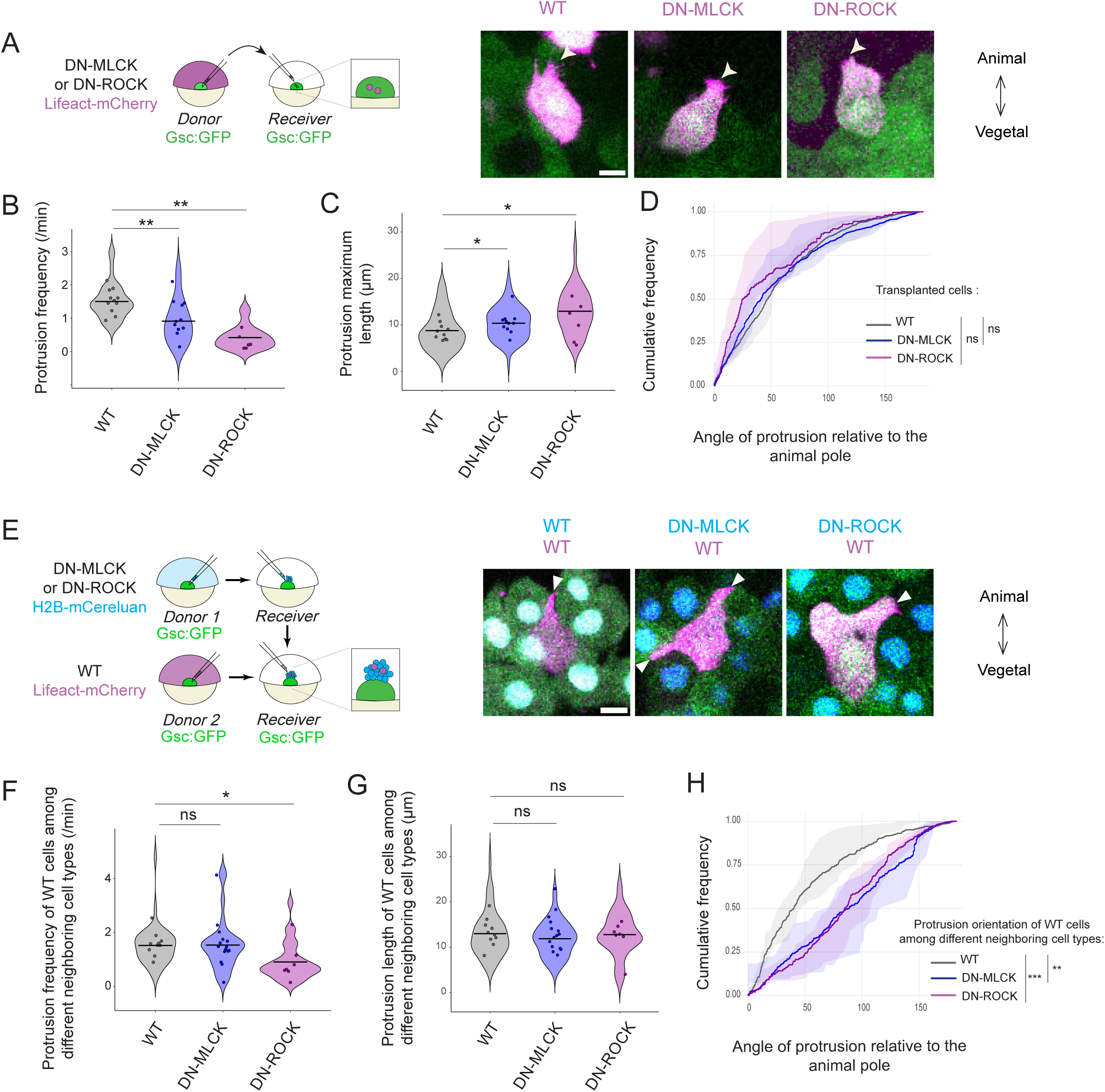
NMII is required in a non-cell autonomous manner to orient polster cells towards the animal pole. (A) Transplants of single cells expressing *Lifeact-mCherry* and *DN-MLCK* or *DN-ROCK* within a wild-type polster in *Tg(-1.8gsc:GFP)* embryos. Scale bar: 10 µm. (B-D) Frequency, maximal length and orientation of protrusions of the transplanted cells (wild-type: 36 cells in 10 embryos, *DN-MLCK*: 32 cells in 11 embryos, *DN-ROCK*: n=23 cells in 6 embryos). Likelihood ratio test of a linear mixed effects model with treatment as a fixed effect and embryos as a random effect against a model without the fixed effect. Adjusted p-values for WT vs *DN-MCLK* and WT vs *DN-ROCK*: (B) 6.0E-3 and 2.2E-3; (C) 0.035 and 0.035; (D) 0.43 and 0.64. (E) Groups of control cells or cells expressing *DN-MLCK* or *DN-ROCK* (and *H2B-Cereluan*) were transplanted at the front of the polster of *Tg(-1.8gsc:GFP)* embryos. Isolated wild-type cells expressing *Lifeact-mCherry* were added into the group. Arrow heads are pointing to cell protrusions. Scale bar: 10 µm. (F-H) Frequency, maximal length and orientation of protrusions of the wild-type transplanted cell, whether it is surrounded by WT, *DN-MLCK* or *DN-ROCK* cells (wild-type: 35 cells in 10 embryos, *DN-MLCK*: 28 cells in 15 embryos, *DN-ROCK*: n=29 cells in 7 embryos). Likelihood ratio test of a linear mixed effects model with treatment as a fixed effect and embryos as a random effect against a model without the fixed effect. Adjusted p-values for WT vs *DN-MCLK* and WT vs *DN-ROCK*: (F) 0.96 and 0.038; (G) 0.74 and 0.74; (H) 5.5E-33 and 1.7E-24.

To directly test this hypothesis, we transplanted a few actin labelled wild-type cells into a small group of NMII inhibited cells, placed at the front of a wild-type polster (Fig. 4E). Inhibiting NMII in neighbouring cells had a very limited effect on the frequency or length of protrusions (Figs. 4F, G). It, however, caused a clear randomization of protrusion orientation (Fig. 4H). These results demonstrate that NMII activity is required within polster cells to control their protrusion dynamics, but is required in neighbouring cells to orient polster cells toward the animal pole.

### NMII is enriched within protrusions and at the rear of cells

In several instances of collective migrations, supracellular actomyosin cables have been observed surrounding the migrating group. These structures contribute to coordinated movement by either restricting cell protrusions at the rear of the group, or by contracting the trailing edge of the group to drive propulsion (Pagès et al., 2022; Reffay et al., 2014; Shellard et al., 2018; Wang et al., 2020). Given the non-cell-autonomous role of NMII in orienting polster cells, we asked whether a similar supracellular cable might be present in the polster. However, NMII signal in *Tg(actb2::myl12.1-gfp)* embryos, as well as P-myosin staining in *Tg(-1.8gsc:GFP)* embryos, revealed no evidence of a cable-like structure surrounding the polster or any supracellular organization of myosin (Figs. 5A, B and Movie S4), suggesting that the non-cell-autonomous function of NMII is mediated through a different mechanism. To get further insight, we looked at the subcellular localization of NMII in migrating polster cells. Actin labelled polster cells from *Tg(actb2::myl12.1-gfp)* embryos were transplanted into the shield of wild-type hosts and imaged during polster migration. Cortical accumulations of NMII were frequently observed, with a pronounced enrichment at the rear of cells (Figs. 5C, D and Movie S5). Notably, these posterior accumulations often coincided with the retraction of the cortex (Fig. 5E), consistent with NMII’s well-established role in driving rear retraction during cell migration (Lauffenburger and Horwitz, 1996). This correlation was confirmed by quantifying cortical dynamics in the presence or absence of NMII accumulations, which showed faster retraction when NMII was enriched (Fig. 5F). In addition to its posterior localization, NMII was also detected within actin-rich protrusions (Fig. 5G and Movie S6). Because obtaining better resolved images of NMII in protrusions proved difficult *in vivo*, we plated *Tg(actb2::myl12.1-gfp)* polster cells on E-cadherin coated coverslips. Under these conditions, polster cells formed small clusters and extended large flat actin-rich protrusions over the coverslip. In this context, NMII was clearly observed accumulating at the base of protrusions (Fig. 5H and Movie S7).

**Figure 5.**
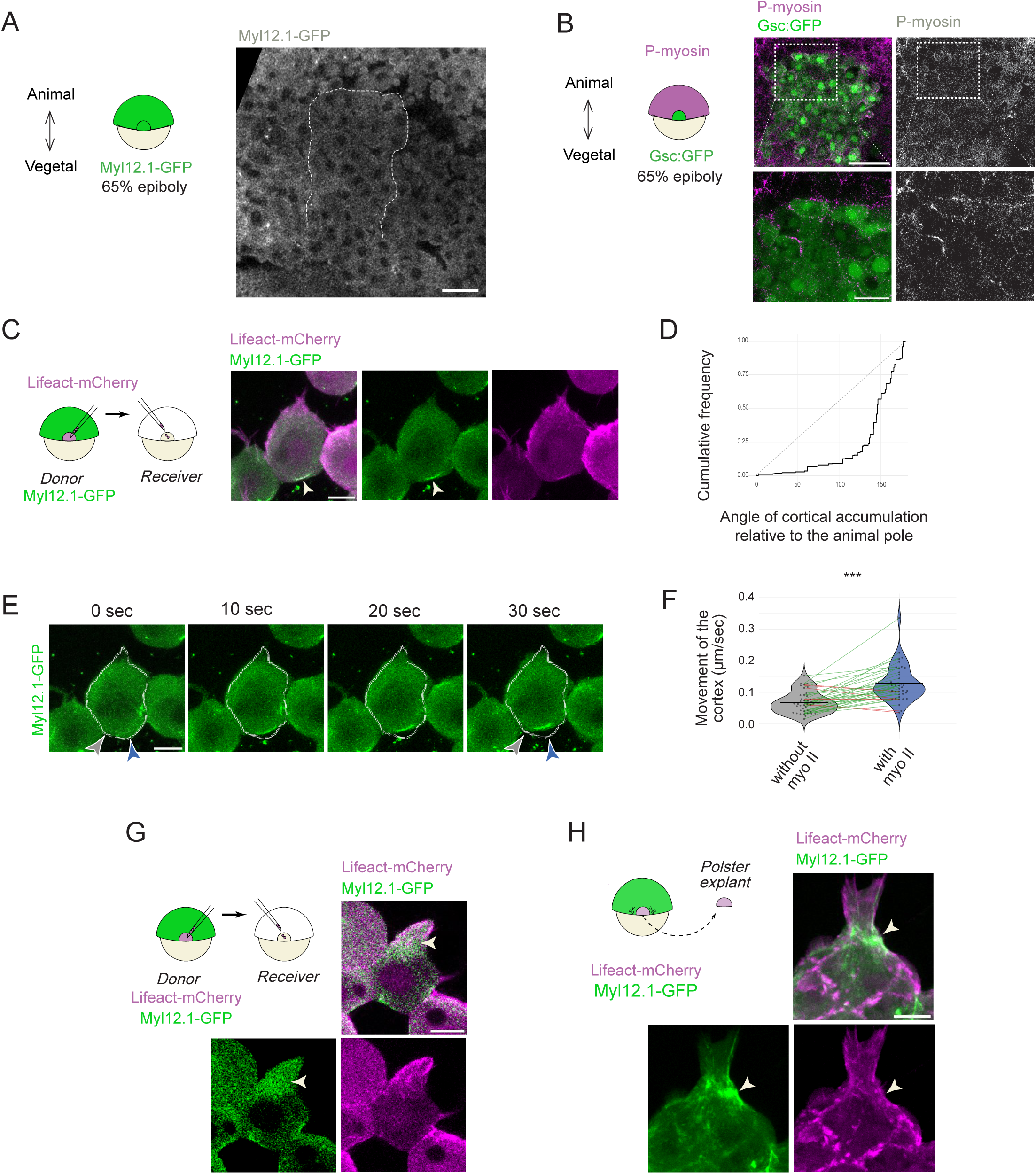
NMII localizes at the posterior cortex and within protrusions of polster cells. (A)NMII signal in *Tg(actb2:myl12.1-EGFP)* embryos at early gastrulation (65% epiboly). Polster cells are recognized as a cluster migrating animalward, delineated by the dashed line (see Movie S4). Scale bar: 30 µm.(B) Immunostaining for phosphorylated myosin light chain type II (P-myosin) in *Tg(-1.8gsc:GFP)* embryos at early gastrulation (65% epiboly). Scale bars: 50 and 20 µm. (C) Transplants of few polster cells expressing *Li-feact-mCherry* from *Tg(actb2:myl12.1-EGFP)* embryos into the polsters of WT embryos. The white arrow head points to a posterior cortical accumulation. Scale bar: 10 µm. (D) Orientation of the cortical accumulations relative to the direction of the animal pole, n = 51 cells in 4 embryos, Kolomogorov-Smirnov uniformity test, p-value: 2.2E-16. (E) Representative example of a polster cell displaying a posterior cortical accumulation of NMII and a concomitant retraction of the cortex (blue arrowhead, compare to grey arrowhead where there is no accumulation of NMII). Scale bar: 10 µm. (F) In 40 cells showing posterior myosin accumulation, cortex movement was measured at the level of the accumulation and at a nearby position without myosin accumulation, 4 embryos, paired t-test, p-value: 1.4E-8. (G-H) Observation of NMII enrichments (arrowheads) in protrusions *in vivo* (G) and *in vitro* (H). n = 26 cells in 4 embryos in vivo, 21 cells in vitro. Scale bar: 10 µm.

### NMII activity controls F-actin retrograde flow and rearward tension in protrusions

At least in some migrating cells, NMII activity contributes to the F-actin retrograde flow observed at the cell leading edge (Lin et al., 2012; Ponti et al., 2004; Qian et al., 2024; Wilson et al., 2010). Given the presence of NMII at the base of polster cell protrusions, we wondered if this could be the case in polster cells. To monitor actin dynamics, we transplanted a few actin labelled polster cells into the polster of unlabelled wild-type hosts, and imaged their protrusions at high temporal resolution. In wild-type cells, we observed a clear retrograde F-actin flow, which seemed to originate from the base of the protrusion (Movie S8). The flow was present both in retracting protrusions and in stable or growing protrusions, suggesting it was then balanced by actin polymerization at the leading edge. Using kymograph analysis, we measured an average retrograde flow speed of 0.16 µm.s^-1^ (Figs. 6A, B). We then analysed cells expressing either *DN-MLCK* or *DN-ROCK* to assess the role of NMII. Inhibiting NMII activity significantly reduced the actin retrograde flow, with average speeds of 0.061 µm.s^-1^ for *DN-MLCK* and 0.078 µm.s^-1^ for *DN-ROCK* (Figs. 6A, B).

**Figure 6.**
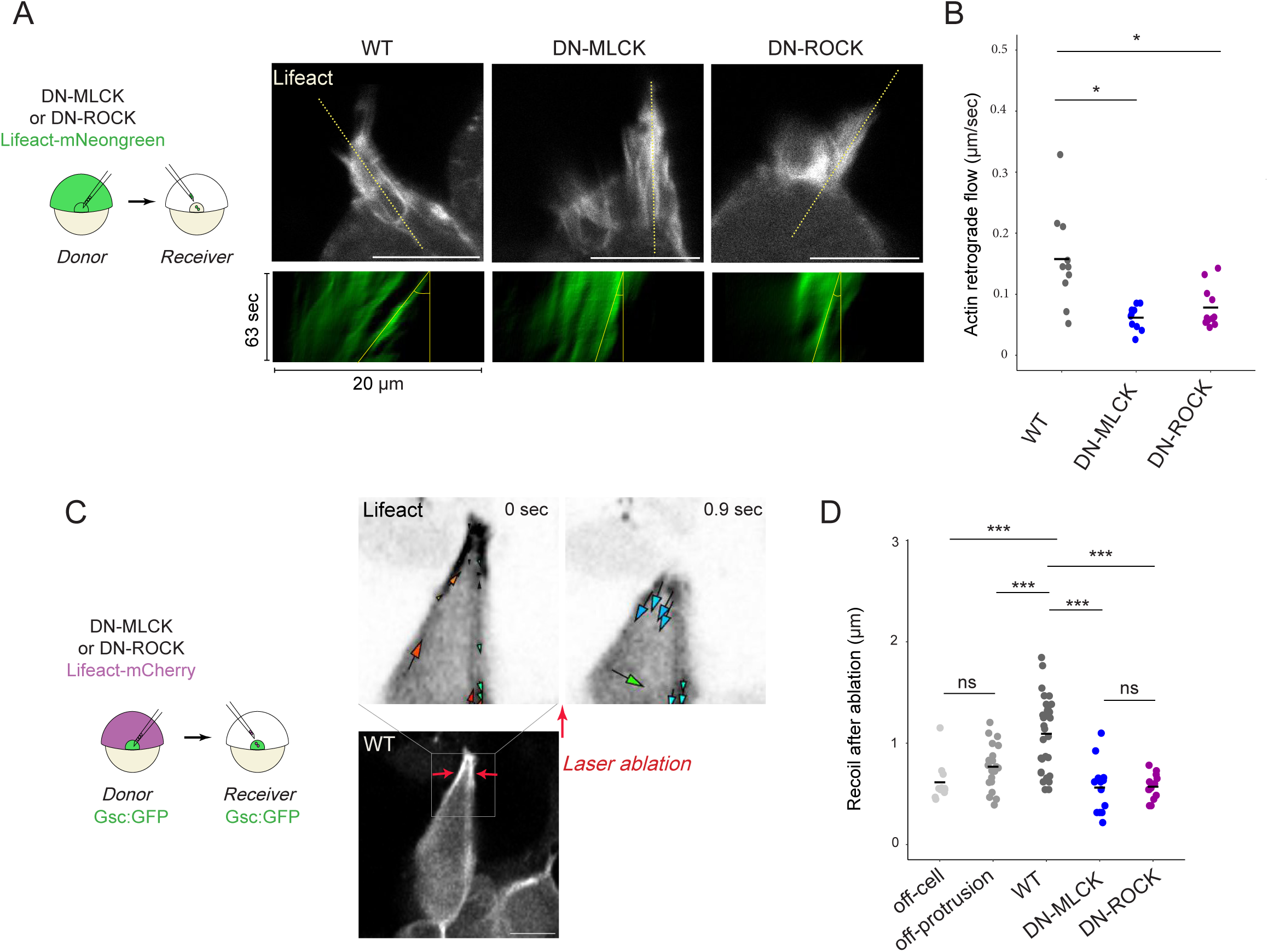
NMII is required in F-actin retrograde flow and protrusion tensioning. (A) Actin-rich protrusions of *Lifeact-mNeongreen* expressing polster cells, transplanted into polsters of wild-type embryos. Kymographs along the yellow dotted line, showing the F-actin retrograde flow (see methods). (B) Actin retrograde flow in wildtype, or *DN-MLCK* or *DN-ROCK* expressing polster cells (n = 10 cells in 4 embryos for WT, 10 cells in 4 embryos for *DN-MLCK*, 11 cells in 4 embryos for *DN-ROCK*). Likelihood ratio test of a linear mixed effects model with treatment as a fixed effect and embryos as a random effect against a model without the fixed effect. Adjusted p-values: 1.5E-3 for WT vs *DN-MCLK*, 4.0E-3 for WT vs *DN-ROCK*. (C) Laser ablations of actin-rich protrusions in Lifeact-mCherry expressing polster cells. The recoil was measured immediately after the ablation. Arrows showing movements were obtained by PIV. (D) Ablations were performed on actin-rich protrusion on wild-type (n = 30), *DN-MLCK* (n = 13) or *DN-ROCK* (n = 13) expressing cells, as well as in regions adjacent to the cell (off-cell; n = 11) and in non-protrusive areas of the cell (off-protrusion; n = 20). One-way ANOVA fol-lowed by Tukey’s HSD post-hoc tests, adjusted p-values: off-cell vs off-protrusion 0.58; off-cell vs WT 4.0E-5; off-protrusion vs WT 9.2E-4; WT vs *DN-MLCK* 1.1E-6; WT vs *DN-ROCK* 1.7E-6; *DN-MLCK* vs *DN-ROCK* 0.99. Scale bars: 10 µm.

According to the molecular clutch model (Elosegui-Artola et al., 2016; Mitchison and Cramer, 1996), retrograde F-actin flow, when coupled to the substrate via adhesions, generates rearward traction forces. We therefore asked whether NMII contributes to tensile forces in polster cell protrusions. To test this, we compared mechanical tensions within the protrusion using recoil after laser ablation as a proxy. Upon ablation of a protrusion, wild-type cells exhibited a limited but significant recoil showing that these protrusions are under tension (Figs. 6C, D and Movie S9). As a control, we either ablated next to a cell without damaging it or targeted the cell cortex in a non-protrusive region; both resulted in significantly smaller recoils compared to protrusion ablations (Fig. 6D). Strikingly, ablation of protrusions in cells expressing *DN-MLCK* or *DN-ROCK* led to a significantly reduced recoil, demonstrating that NMII is required to generate tensile forces within polster cell protrusions (Fig. 6D).

### NMII is required non-cell-autonomously to open α-catenin

We initially set out to investigate the role of NMII in polster cell migration based on our previous finding that the mechanosensitive M-domain of α-catenin is required for polster cells orientation. In epithelial cells, the opening of this domain is triggered by NMII activity pulling on the adherens junction complex. We hypothesized that NMII might serve a similar function in polster cells. The open conformation of α-catenin’s M-domain can be detected using the α18 antibody (Yonemura et al., 2010). We immunostained embryos expressing *β-gal, DN-MLCK,* or *DN-ROCK* for both total α-catenin and α18 (Fig. 7A). Inhibition of NMII induced a reduction in the α18/α-catenin ratio, consistent with a role of NMII in α-catenin opening (Fig. 7B).

**Figure 7.**
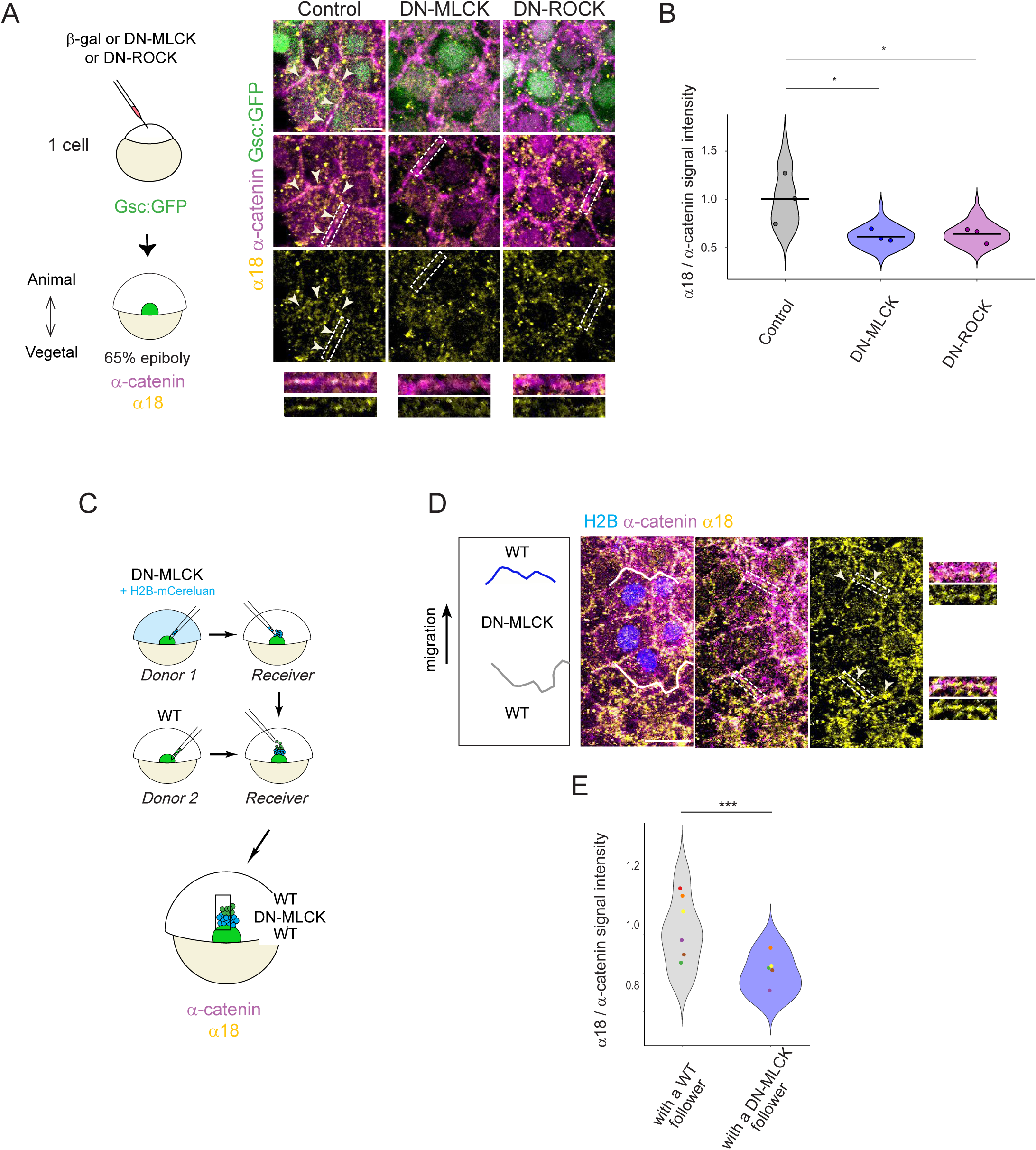
NMII is required in a non-cell autonomous manner to open the mechano-sensitive domain of α-catenin. (A) Immunolabelling of α-catenin and of the open form of α-catenin (α18 antibody) in *Tg(-1.8gsc:GFP)* embryos expressing *β-gal*, *DN-MLCK* or *DN-ROCK*. Scale bar: 10 µm. (B) Ratio of α18 and α-catenin intensities at the cortex of polster cells (303 cell-cell junctions for control, 222 for *DN-MLCK*, 178 for *DN-ROCK*; 3 embryos per condition). Likelihood ratio test of a linear mixed effects model with treatment as a fixed effect and embryos as a random effect against a model without the fixed effect. Adjusted p-values: Ctl vs *DN-MCLK* 0.022; Ctl vs *DN-ROCK* 0.022. (C-D) Wildtype cells were transplanted ahead of *DN-MLCK* expressing cells, placed at the front of the polster of a wildtype *Tg(-1.8gsc:GFP)* embryo. Immunolabelling of α-catenin and of the open form of α-catenin (α18 antibody). (E) Ratio of α18 and α-catenin intensities at the cell-cell junction between a WT polster cell followed by a *DN-MLCK* polster cell, or a *DN-MLCK* polster cell followed by a WT cell. n = 6 embryos, 20 junctions with a WT cell followed by a *DN-MLCK* cell, 19 with *DN-MLCK* followed by WT. Likelihood ratio test of a linear mixed effects model with type of cell junction as a fixed effect and embryos as a random effect against a model without the fixed effect. p-value: 2.4E-5. Dots are mean per embryo, one colour per embryo.

However, our study also revealed that NMII is not required cell-autonomously for polster cell orientation. Instead, it acts non-cell-autonomously. Moreover, we observed NMII enrichment at the base of cellular protrusions, and showed that it is necessary to generate rearward tension in these structures. Altogether, these observations point to a model in which the tension required to open α-catenin in a given cell is not generated by that cell’s own NMII activity, but rather by NMII activity in the protrusion of the following cell. To directly test this hypothesis, we transplanted a few wild-type cells in front of a small group of NMII-inhibited cells, positioned at the front of a wild-type polster (Fig. 7C). We then quantified the α18/α-catenin ratio at cell-cell junctions between wild-type cells followed by NMII inhibited cells, and NMII inhibited cells followed by wild-type cells (Fig. 7D-E). The ratio was significantly lower when the following cell is NMII-inhibited, demonstrating that NMII is required in a non-cell autonomous manner in follower cells to induce the conformational opening of α-catenin’s mechano-sensitive domain in leading cells.

## Discussion

Our study reveals an unexpected non-cell-autonomous role for non-muscle myosin II (NMII) in guiding collective cell migration. Using a combination of genetic, pharmacological, and mosaic transplantation approaches in the zebrafish polster, we show that NMII activity is required not only for cell-autonomous functions (F-actin retrograde flow and protrusion dynamics) but also for coordinating the orientation of neighbouring cells via intercellular tension. We demonstrate that NMII localizes to the rear cortex and to the base of protrusions, where it generates rearward traction forces. These forces are transmitted through cell–cell contacts to mechanically open the M-domain of α-catenin in adjacent cells. This intercellular force transmission orients neighbouring cells towards the animal pole, thereby coordinating the collective migration of the polster.

We previously established that the mechanosensitive domain of α-catenin is required for cell-cell contacts to orient cell migration, likely via vinculin recruitment, PI3K activation (Boutillon et al., 2022), and yet-to-be-identified downstream effectors, potentially including Merlin (Das et al., 2015). In the present study, we aimed to identify the upstream event that triggers α-catenin activation (likely opening), as this represents the signal transmitted between cells to coordinate migration. In epithelial cells, α-catenin opening during adherens junction biogenesis is induced by NMII contraction of the branched actin network at the junction (Heuzé et al., 2019). More recently, in migrating epithelial cells, antiparallel cortical F-actin flows at lamellipodia during cell-cell collisions—rather than myosin activity—were proposed to open α-catenin and mediate contact inhibition of locomotion (Noordstra et al., 2023). In contrast, our findings in the mesenchymal polster cells reveal that α-catenin opening is triggered by an externally applied force: NMII-dependent rearward tension exerted by protrusions from neighbouring cells. This observation raises one question: for α-catenin to be opened by an external force, the force must be opposed by anchoring structures that allow tension to build. We observed NMII accumulations at the rear of polster cells that could generate such resistance. However, NMII-inhibited cells still exhibited normal orientation and α-catenin opening, suggesting that NMII is not essential for resisting the protrusive traction. An alternative is that α-catenin at the cell rear is anchored to a sufficiently stiff actin cortex that passively resists tension. Notably, during imaging actin flow within protrusions, we identified protrusions likely engaging neighbouring cells. In these cases, retrograde actin flow in the protrusion often correlated with deformation of the neighbouring cell at the contact site, and, in some instances, with cortical actin buckling (Movie S10). These observations confirm that protrusions exert tension on neighbouring cells and suggest that adherens junctions are linked to the cortical actin network and under mechanical tension.

### Protrusions as conveyors of directional information

Protrusions are traditionally viewed as grappling hooks: they extend through actin polymerization, grip the substrate, and are used to pull the cell forward (Fulga and Rørth, 2002). Such a role was recently nicely demonstrated *in vivo*, in the migrating primordium of the fish lateral line, where NMII localizes at the base of protrusions and drives rearward F-actin flow to propel the primordium (Qian et al., 2024). NMII accumulations at protrusion bases have also been reported in Drosophila border cells. However, in this system, detailed analysis of cell body and protrusion dynamics revealed that protrusion retraction often does not correlate with cell body advancement, challenging the simple “grapple-and-pull” model. Instead, protrusions have been proposed to serve as sensory organs, probing the environment for physical openings in which the cells can then migrate (Mishra et al., 2019). In the polster, we observed NMII at the base of cell protrusions, where it promotes F-actin flow and rearward tension. While these protrusions may indeed act as grapples or sensors, our findings uncover an additional, instructive role. The rearward tension generated by NMII in a protrusion serves to mechanically open α-catenin in the neighbouring cell, transmitting directional information. This new function resolves a longstanding puzzling observation: why are all cells within the polster protrusive, even central cells, completely surrounded by other polster cells? Under a simple “grapple-and-pull” hypothesis, peripheral cells extend protrusions to pull on the substrate, but the usefulness of protrusions in central cells is not obvious —any pull they exert on neighbours would propel them but pull the neighbours backward, resulting in no net group movement. Instead, if each cell’s protrusion transmits an orienting force to its immediate neighbour, then every cell, regardless of position, contributes to a relay of directional cues. In this view, central cells are not misguidedly tugging on their neighbours; rather, they transmit an information that aligns the entire group.

### NMII and supracellular actomyosin in collective migration

Our results reveal a non-cell-autonomous role for NMII in coordinating a collective migration. NMII non-cell-autonomous roles have been reported in other collective migrations, including epithelial sheet closure (Reffay et al., 2014), border cell migration in Drosophila (Mishra et al., 2019), neural crest cell migration (Shellard et al., 2018), or migration of cancer cell clusters (Pagès et al., 2022). In these contexts, however, NMII acts as part of a supracellular actomyosin cable that encircles the cell cohort, and serve to either restrict protrusive activity at the rear (Reffay et al., 2014; Wang et al., 2020), contract at the trailing edge to drive cellular treadmilling (Shellard et al., 2018), or generate oscillatory deformations that propel the cohort (Pagès et al., 2022). These cables operate at a tissue scale, shaping the group’s polarity and propulsion. In the polster, however, we found no evidence of a circumferential myosin cable. Instead, NMII acts locally—at each cell’s rear cortex and protrusion base—to generate forces that propagate from cell-to-cell.

Force propagation across cell-cell contacts has been proposed to polarize cells across epithelial monolayers in culture, and, *in vivo*, in Drosophila border cells (Cai et al., 2014). In these systems, force generation in mainly achieved by leader cells and transmitted to followers to synchronize motion. However, accumulating evidence shows active roles of follower cells in generating forces and influencing leader cells, suggesting the mechanisms observed in this study may apply more broadly to other systems (Farooqui and Fenteany, 2005; Tambe et al., 2011; Vishwakarma et al., 2018). Of particular interest is whether similar mechanisms operate in cancer cell migrations, many of which are collective and dependent on NMII (Doran et al., 2024; Halder et al., 2021; Pagès et al., 2022).

## Acknowledgements

We thank Emilie Menant for fish care; P. Mahou and the Polytechnique Bioimaging Facility for imaging on their equipment supported by Region Ile-de-France (interDIM) and Agence Nationale de la Recherche (ANR-11-EQPX-0029 Morphoscope2, ANR-24-INBS-0005 France-BioImaging BIOGEN). This work was supported by the ANR grants ANR-20-CE13-0016-03 and ANR-25-CE13-6448-01, the Serge Schoen New Synergies Grant Program from the Engineering for Health center. AE was supported by Fondation pour la Recherche Médicale grant FDT202404018639.

## Author Contributions

AE, AB and ND conceived experiments, which were performed by AE and AB. AE analyzed data. AE and ND wrote the manuscript. ND secured funding.

## Materials & Methods

### Key resources table

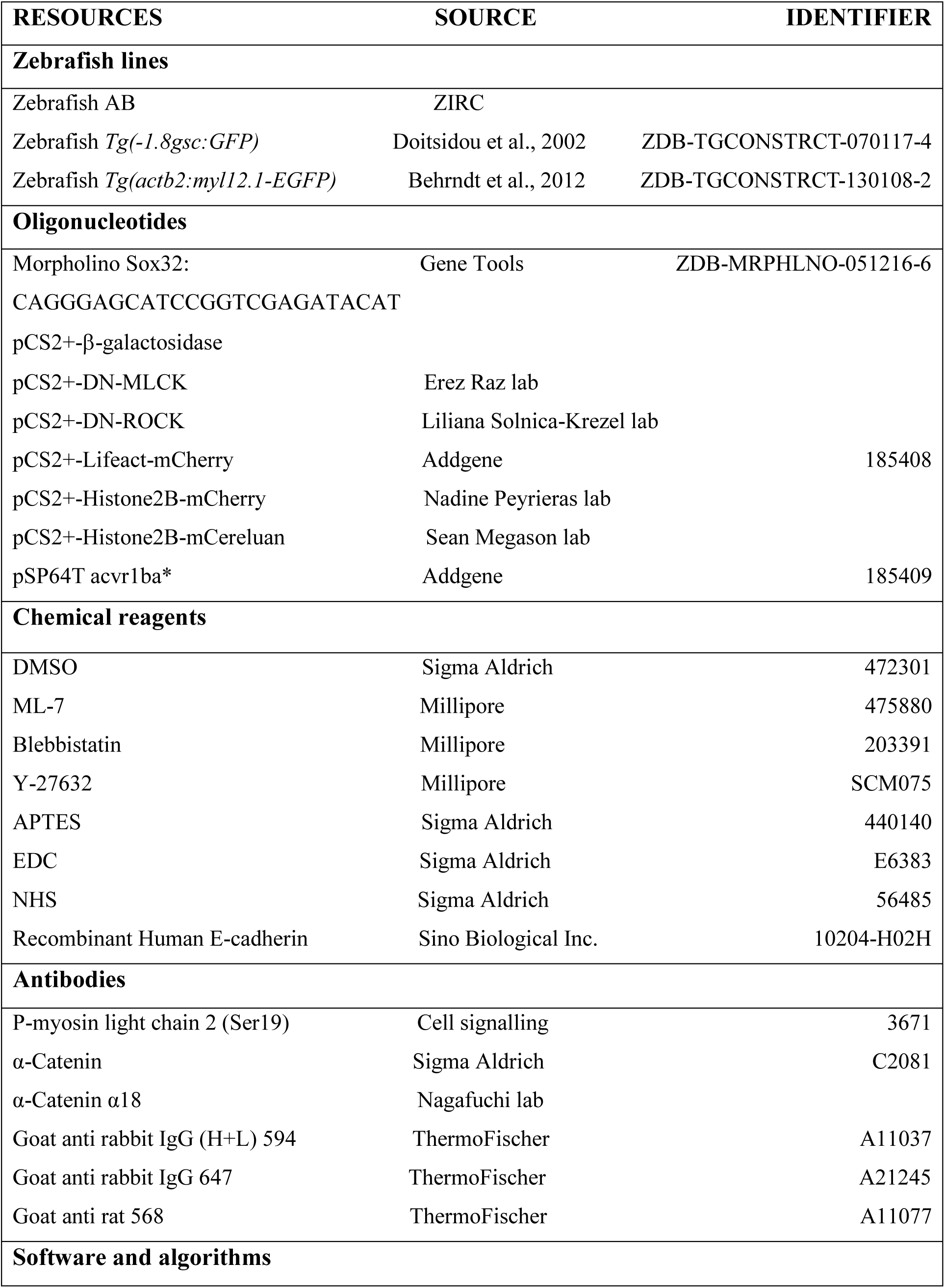
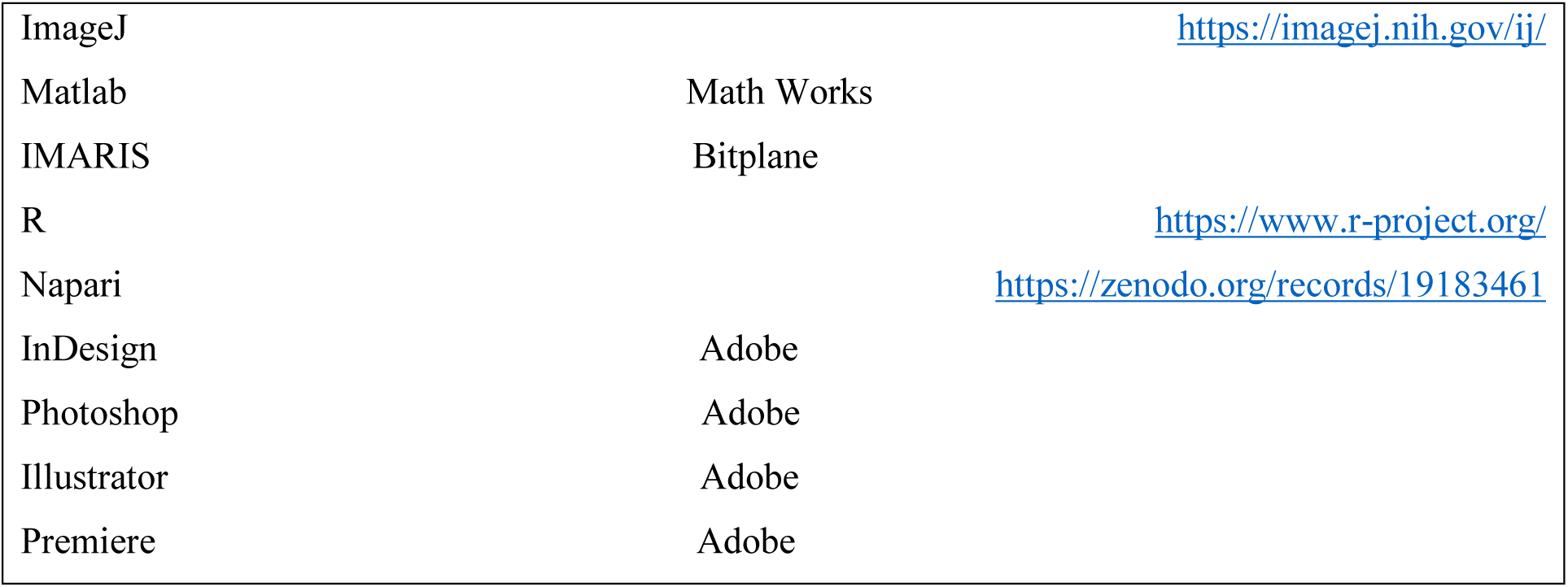

### Zebrafish lines and husbandry

Embryos were obtained by natural spawning of AB (wild-type), *Tg(-1.8gsc:GFP)* and *Tg(actb2:myl12.1-EGFP)* adult fishes (Behrndt et al., 2012; Doitsidou et al., 2002). *Tg(-1.8gsc:GFP)* in which GFP is expressed under the control of *goosecoid* regulatory sequences (*gsc*) allows identification of the polster. GFP is also expressed in the following axial mesodermal cells as well as in some endodermal cells. *Tg(actb2:myl12.1-EGFP)* expresses ubiquitously a GFP-tagged form of the NMlI regulatory light chain. Embryos were incubated at 28°C and staged in hours post-fertilization (hpf) (Kimmel et al., 1995). All animal experimentation was conducted in accordance with the local ethics committee and approved by the Ethical Committee N°59 and the Ministère de l’Education Nationale, de l’Enseignement Supérieur et de la Recherche under the file number APAFIS #44004-2023062815026564 v2.

### Zebrafish cell injections

Capped sense mRNAs were synthesized from pCS2+ or pSP64T constructs with the mMessage mMachine SP6 kit (Thermo Fischer) and used at the following concentrations: *acvr1ba\** (0.6 ng/µL), *DN-MLCK* (50 ng/µL), *DN-ROCK* (5 ng/µL when injected at the one-cell stage, 2.5 ng/µL when injected at the four-cell stage), *β-galactosidase* (50 ng/µL), *Lifeact-mCherry* (30 ng/µL), *Lifeact-mNeongreen (30 ng/µL), H2B-mCherry* (30 ng/µL), *H2B-mCereluan* (30 ng/µL). Global inhibition of NMII from the one-cell stage was achieved by injecting *DN-MLCK* or *DN-ROCK* mRNAs at the 1-cell stage in *Tg(-1.8gsc:GFP)* embryos, together with *H2B-mCherry* to visualize nuclei and track polster cells. *β-galactosidase (β-gal)* was used as control mRNA. At 60%-epiboly (6.5 hpf), embryos were selected and mounted for imaging. Inhibition of NMII during gastrulation was performed in *Tg(-1.8gsc:GFP)* embryos, injected with *H2B-mCherry* at the 1-cell stage. At the germ-ring stage (5 hpf), embryos were treated with ML-7 (25 µM) or blebbistatin (50 µM), both dissolved in DMSO, or Y-27632 (20 µM) dissolved in water, until 60%-epiboly (6.5 hpf). Embryos were then mounted for imaging in the presence of the NMII inhibitors. To impose a polster identity in transplant experiments, *acvr1ba\** mRNA and *sox32* morpholino (5’-CAGGGAGCATCCGGTCGAGATACAT-3’; Gene Tools LLC Philomath; 0.3 mM) were injected into one cell at the 4-cell stage (Dumortier et al., 2012).

### Cell transplants and microsurgery

Experiments to inhibit myosin activity (polster replacement, cell autonomous and non-cell autonomous inhibition) were performed in *Tg(-1.8gsc:GFP).* Polster removals were performed at 60%-epiboly (6.5 hpf) by aspiration with a large homemade glass pipette. Around 70 to 100 polster cells expressing *β-gal* or *DN-MLCK* or *DN-ROCK* with *H2B-mCherry* were then transplanted into these embryos, ahead of the notochord. Embryos were then selected and mounted for imaging around 65%-epiboly (7 hpf). For cell-autonomous inhibition of NMII, individual polster cells expressing *DN-MLCK* or *DN-ROCK* together with *Lifeact-mCherry* were transplanted into the shield of receiver embryos at shield stage (6 hpf). For non-cell autonomous inhibition of NMII, clusters of 10-20 polster cells expressing *DN-MLCK* or *DN-ROCK* and labelled with *H2B-mCereluan* were transplanted to host embryos at shield stage (6 hpf). Single or few wild-type polster cells expressing *Lifeact-mCherry* were transplanted into these clusters. Embryos were then selected and mounted for imaging around 65%-epiboly (7 hpf).

To analyse the intracellular localization of NMII, polster cells from *Tg(actb2:myl12.1-EGFP)* embryos expressing *Lifeact-mCherry* were transplanted into the shield of AB receiver embryos. Embryos were mounted for imaging at 60%-epiboly (6.5 hpf).

### Polster cell culture

At 60%-epiboly (6.5 hpf), the polsters of *Lifeact-mCherry* expressing *Tg(actb2:myl12.1-EGFP)* embryos were manually dissected and placed in Ringer’s without calcium, to induce cell dispersion. Dispersed cells were then transferred into a drop of culture medium on an E-cadherin coated glass coverslip. Briefly, for coating, coverslips were silanized with APTES, before incubation with E-cadherin previously mixed to EDC and NHS. Cells were incubated at 28°C for 30 minutes to allow cells to attach to the substrate and imaged using an inverted SP8 (Leica) confocal microscope. Culture Medium: 80% Leibovitz’s L-15 medium (Thermofischer) diluted at 65% in water + 20% Embryo medium + 0.1% BSA + 125 mM HEPES (ph7.5) + 10U/mL Penicillin/Streptomycin.

### Immunostaining

Immunostainings were performed on *Tg(-1.8gsc:GFP)* embryos expressing *β-gal* or *DN-MLCK* or *DN-ROCK.* Embryos were fixed overnight in 4% PFA (pH 7) at 4°C. They were then rinsed with PBT (PBS 1X + 0.1% Tween20), dechorionated, and incubated for one hour in 5% goat serum prepared in PBDT (PBS 1X + 1% BSA + 1% DMSO + 0.1% Tween20). The primary antibodies and dilutions used were: P-myosin (rabbit) (1:200), α-Catenin (rabbit) (1:200) and α18 (rat) (1:200). Embryos were washed four times in PBDT. Secondary antibodies were applied at the following dilutions: goat anti-rabbit IgG (H+L) 594 (1:1000), goat anti-rabbit IgG 647 (1:700) and goat anti-rat 568 (1:100). Embryos were then washed four additional times in PBDT before mounting.

### Imaging

Whole embryo imaging was performed with a M205FCA stereomicroscope (Leica) equipped with a MC170HD Camera (Leica). Imaging of embryos for analysing protrusion dynamics, immunolabelling and the NMII subcellular localization were carried out using a Leica TCS SP8 inverted confocal microscope (Leica), equipped with a 28°C chamber with a HC PL APO 40x/1.10WCS2 objective (Leica). For cell tracking and imaging of *Tg(actb2:myl12.1-EGFP)* embryos, we used a TriM Scope II two-photon microscope (La Vision Biotech) equipped with a 28°C chamber (okolab) and an XLPLN25XWMP2 (Olympus) 25x water immersion objective. Movies were acquired at a time interval of two minutes and with a Z-step of 2 microns. The MaiTai laser was set to 920 nm with a power of 15% to image either polster cells or the NMII signal, and the Insight laser was set to 1160 nm with a power of 15% to image the nuclei.

### Laser ablations

Protrusion ablations were performed under The TriM Scope II two-photon microscope on *Lifeact-mCherry* labelled cells. mCherry was imaged with the Insight laser at 1160 nm, while ablations were performed with the Mai Tai laser at 820 nm and an exit power of 0.3 mW at the lens. To minimize the time interval between frames to 0.3 s, imaging was restricted to 50 µm wide regions. Ten images were recorded before ablation. Ablation was performed on a 3*7 microns area across the protrusion. Imaging resumed immediately after ablation, and continued for 50 time frames. Recoil was measured both manually in ImageJ and by PIV analysis in Matlab (Math Works).

### Image analyses

Leading edge progression, polster width and polster distance to the margin were quantified in ImageJ. Labelled nuclei were tracked in Imaris (Bitplane) and further processed using Matlab (Math Works) to compute instantaneous speed, speed in the direction of the animal pole (axial velocity) and cell orientation (defined as the angle of each cell’s displacement relative to the direction of the animal pole). Instantaneous speed was determined as the norm of the cell movement divided by the time interval between two acquisitions (two minutes). Axial velocity was calculated as the norm of the displacement vector projected onto the vegetal-animal axis, divided by the time interval. For *Tg(actb2:myl12.1-EGFP)* embryos, images were processed in Imaris, where the polster was identified as the anterior most cluster of cells migrating towards the animal pole throughout the time-lapse. Analyses of immunolabelled embryos were performed in ImageJ and Napari. Polster cells were identified by expression of GFP (*gsc:GFP* line). Cell-cell junctions were manually segmented, and P-myosin, α-catenin, and α18 mean intensities were measured over a 2-micron-wide region along the junction. Actin-rich protrusions were quantified in cells expressing *Lifeact-mCherry* or *Lifeact-mNeongreen* using ImageJ. Frequency was quantified as the number of protrusions per cell observed at each time point throughout the entire movie. Protrusion length was determined by measuring the maximal length of a given protrusion. The orientations of protrusions or cortical accumulations of NMII were measured relative to the direction of the animal pole. Retrograde actin flow was assessed by generating kymographs over a 2.8 µm wide (50-pixel) band along the protrusion with the KymographClear plugin (Mangeol et al., 2016). This plugin separates backward and forward motions, through a Fourier transform based algorithm. Measurements of cortex speed with and without NMII accumulations were performed manually in ImageJ by measuring the cortex displacement between two frames and dividing it by the time interval.

### Illustrations

Figures were assembled with Adobe InDesign, schematics were created in Adobe Illustrator. Images of embryos were processed in Adobe Photoshop. Movies were assembled in Adobe Premiere.

### Statistics

Plots and statistical analyses were performed in R (R Core Team, 2024). For independent observations, significance was calculated using t-tests to compare means between two experimental conditions or a one-way ANOVA followed by Tukey’s HSD post-hoc test for comparisons involving more than two conditions. For non-independent observations (several cells for each embryo), we used lme4 (Bates et al., 2015), to perform mixed effects analyses. As fixed effects we entered the treatment (for instance Control, DN-MLCK, DN-ROCK), as random effects we had intercepts for embryos. p-values were obtained by likelihood ratio tests of the full model with the fixed effect against the model without the fixed effect. When pairwise comparisons were performed, p-values were adjusted using the Benjamini Hochberg method. In all cases, normality and homoscedasticity were verified by visual inspection of residual plots and through Shapiro and Levene tests. Sample size, used tests and p-values are indicated in the figure legends. In all figures, ns: p-value ≥0.05; *: p<0.05; **: p<0.01; ***: p<0.001.

---

**Supplementary Figure 1.**
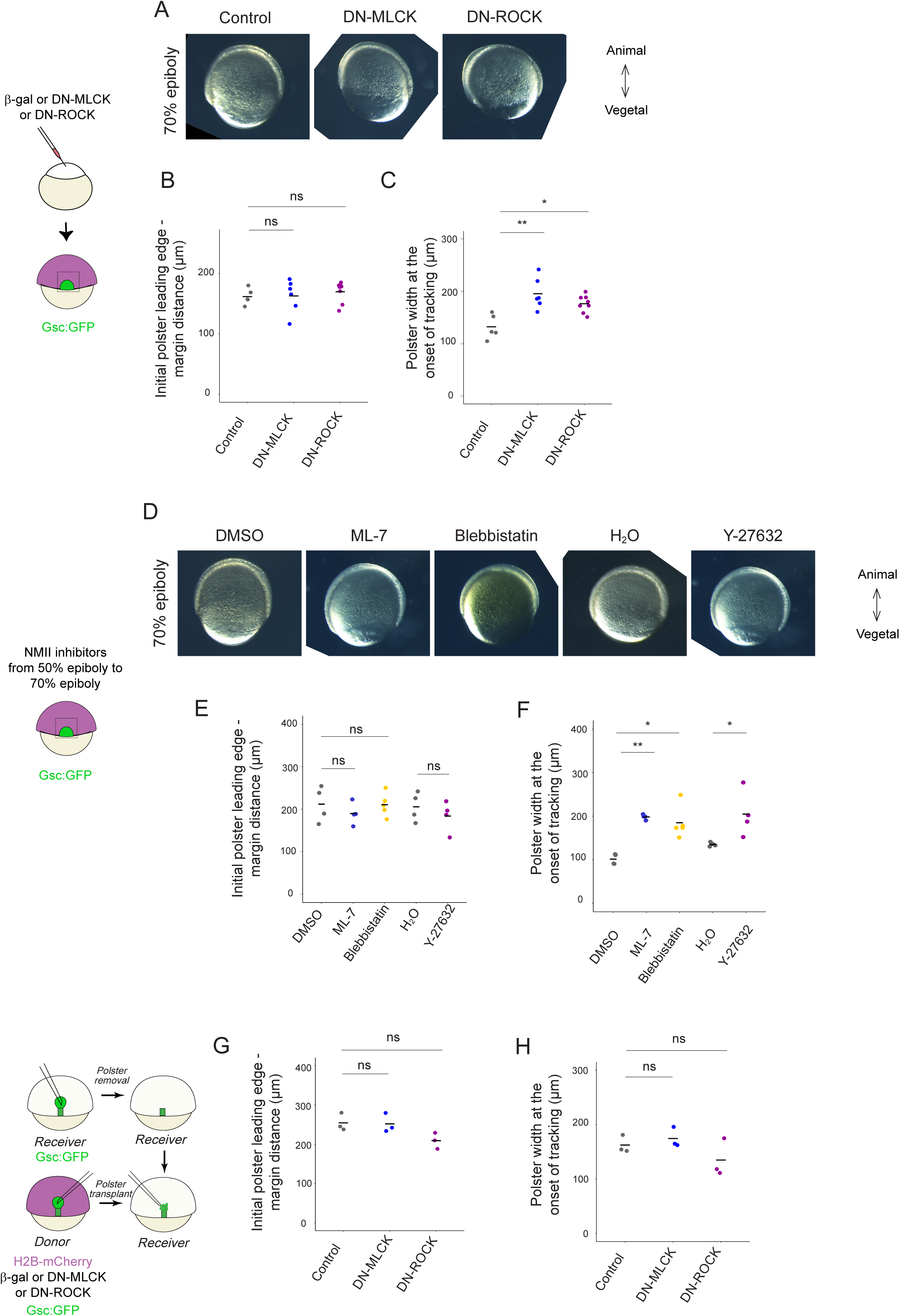
Characterisation of the polster at the onset of tracking. (A) Embryos expressing *β*-*galactosidase* (control), *DN-MLCK* or *DN-ROCK* at mid-gastrulation, after polster tracking. (B) Distance between the leading-edge of the polster and the margin, and (C) width of the polster, at the onset of polster tracking (60% epiboly). n = 5, 6 and 8 embryos for *β*-*gal*, *DN-MLCK* and *DN-ROCK*. For edge to margin distance, one-way ANOVA, p-value = 0.73. For polster width, one-way ANOVA followed by Tukey’s HSD post-hoc tests, adjusted p-values: Control vs *DN-MLCK*: 8.9E-4; Control vs *DN-ROCK*: 0.010. (D) Embryos at mid-gastrulation, treated with NMII inhibitors starting shortly before the onset of gastrulation (germ ring, 5 hpf). (E) Distance between the leading-edge of the polster and the margin, and (F) width of the polster, at the onset of polster tracking (60% epiboly). n = 4, 5, 4, 4 and 4 embryos for DMSO, ML-7, blebbistatin, H2O and Y-27632. t-test or one-way ANOVA, followed by Tukey’s HSD post-hoc tests when significant. For edge to margin distance, for DMSO, ML-7 and blebbistatin, p-value = 0.55; for H2O vs Y-27632: 0.42. For polster width, 6.4E-4 for DMSO vs ML-7, 1.3E-3 for DMSO vs blebbistatin, 0.077 for H2O vs Y-27632. (G) Distance between the leading-edge of the polster and the margin, and (H) width of the polster, at the onset of polster tracking (65% epiboly), in embryos where the endogenous polster was replaced with a polster expressing *β*-*gal* or *DN-MLCK* or *DN-ROCK*. n = 3 embryos for each condition. One-way ANOVA: p-value = 0.08 for edge to margin distance; 0.21 for polster width.

**Supplementary Figure 2.**
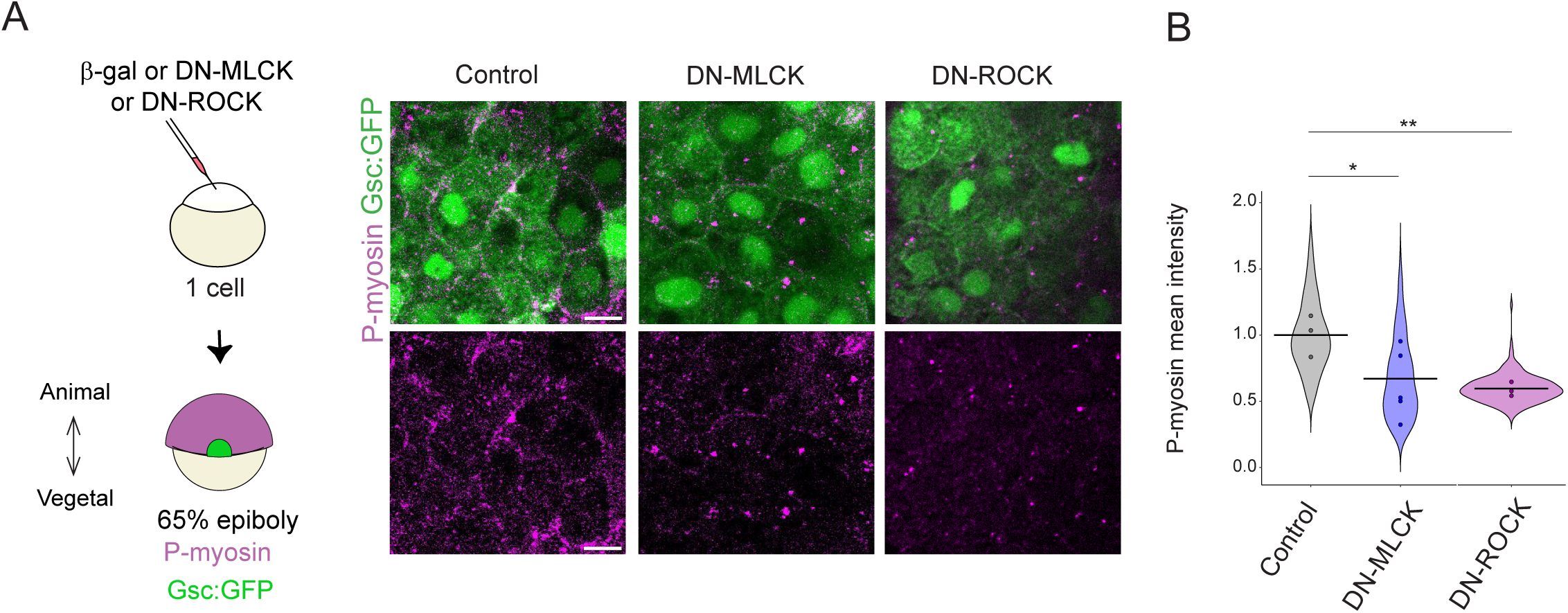
DN-MLCK and DN-ROCK reduce phosphorylated NMII in polster cells. (A) Immunostaining for phosphorylated myosin light chain type II (P-myosin) in *Tg(-1.8gsc:GFP)* embryos expressing *β-gal* or *DN-MLCK* or *DN-ROCK*, fixed at early gastrulation (65% epiboly). (B) Quantification of P-myosin mean intensity at the cortex of polster cells (377 cell-cell junctions in 3 embryos for control, 544 in 5 embryos for *DN-MLCK*, 192 in 3 embryos for *DN-ROCK*). Likelihood ratio test of a linear mixed effects model with treatment as a fixed effect and embryos as a random effect against a model without the fixed effect. Adjusted p-values: Ctl vs *DN-MCLK* 0.028; Ctl vs *DN-ROCK* 0.0026

## Movie legends

**Movie S1. Polster migration in NMII inhibited embryos.**

Time-lapse imaging of *Tg(-1.8gsc:GFP)* embryos expressing *β-gal* (Control) or *DN-MLCK* or *DN-ROCK*, showing leading-edge progression over a 40-minute period, and the trajectories of polster cells.

**Movie S2. NMII is required during gastrulation for polster cell migration.**

Time-lapse imaging of *Tg(-1.8gsc:GFP)* embryos treated with different NMII drug inhibitors, starting shortly before the onset of gastrulation, showing leading-edge progression over a 40-minute period, and the trajectories of polster cells.

**Movie S3. NMII is specifically required in the polster for its orientation towards the animal pole.**

Time-lapse imaging of *Tg(-1.8gsc:GFP)* embryos in which the endogenous polster was replaced with polster cells expressing *β-gal* (Control) or *DN-MLCK* or *DN-ROCK*, showing leading-edge progression over a 40-minute period, and the trajectories of polster cells.

**Movie S4. NMII is specifically required in the polster for its orientation towards the animal pole.**

4D imaging of *Tg(actb2:myl12.1-EGFP)* embryos at early gastrulation. The movie first shows the signal across a z-stack, and then shows how the leading-edge of the polster (red arrow head) can be identified as the edge on an animalward-migrating cell cluster. No supracellular myosin structures are observed within the polster.

**Movie S5. NMII accumulation at the rear of a polster cell.**

Time-lapse imaging of a *Tg(actb2:myl12.1-EGFP)* polster cell expressing *Lifeact-mCherry*.

The white arrowhead points to a posterior cortical accumulation of NMII.

**Movie S6. NMII accumulation in a protrusion.**

Time-lapse imaging of a *Tg(actb2:myl12.1-EGFP)* polster cell expressing *Lifeact-mCherry*.

NMII accumulation is visible in a protrusion.

**Movie S7. NMII accumulation at the basis of protrusions.**

3D rendering of *Tg(actb2:myl12.1-EGFP)* polster cells expressing *Lifeact-mCherry,* plated on an E-cadherin coated coverslip. Arrow heads point to NMII accumulations at the basis of protrusions.

**Movie S8. Actin retrograde flow in polster cell protrusions.**

High temporal resolution time-lapse showing the actin retrograde flow (Lifeact-mNeongreen labelling), in protrusions of control or *DN-MLCK* or *DN-ROCK* expressing polster cells, transplanted into the polster of unlabelled wild-type host embryos.

**Movie S9. Laser ablation of protrusions.**

Ablations were performed on actin-rich protrusion on wild-type, DN-MLCK or DN-ROCK expressing cells, as well as in regions adjacent to the cell and in non-protrusive areas of the cell.

**Movie S10. Laser ablation of protrusions.**

High temporal resolution time-lapse of *Lifeact-mNeongreen* expressing polster cells, showing a protrusion engaging a neighbouring cell. Retrograde actin flow in the protrusion correlates with local deformation and cortical actin buckling (arrow heads) in the neighbouring cell.

